# Illuminating chromatin compaction in live cells and fixed tissues using SiR-DNA fluorescence lifetime

**DOI:** 10.1101/2020.05.02.073536

**Authors:** Colin Hockings, Chetan Poudel, Kevin A. Feeney, Clara L. Novo, Mehdi S. Hamouda, Ioanna Mela, David Fernandez-Antoran, Pedro P. Vallejo-Ramirez, Peter J. Rugg-Gunn, Kevin Chalut, Clemens F. Kaminski, Gabriele Kaminski-Schierle

## Abstract

The global compaction state of chromatin in a nucleus is an important component of cell identity that has been difficult to measure. We have developed a quantitative method to measure the chromatin compaction state in both live and fixed cells, without the need for genetic modification, using the fluorescence lifetime of SiR-DNA dye. After optimising this method using live cancer cell lines treated to induce chromatin compaction or decompaction, we observed chromatin compaction in differentiating epithelial cells in fixed tissue sections, as well as local decompaction foci that may represent transcription factories. In addition, we shed new light on chromatin decompaction during embryonic stem cell transition out of their naïve pluripotent state. This method will be useful to studies of nuclear architecture, and may be easy, cheap, and accessible enough to serve as a general assay of ‘stem-ness’.

## Introduction

Chromatin architecture plays a crucial role during differentiation and in the maintenance of cell identity. However, a quick and accessible method to determine the level of chromatin compaction or decompaction is lacking. Currently, most studies of nuclear architecture rely on genomic sequencing-based methods to measure chromatin compaction, such as High Throughput Chromosome Conformation Capture (Hi-C) (1–4), Chromatin Immunoprecipitation Sequencing (ChIP-seq) (5, 6) or Assay for Transposase-Accessible Chromatin using sequencing (ATAC-seq) (7). These methods produce outstanding detail on local chromatin structure, domain structure and long-range interactions, but generally represent an expensive snapshot of a populations of cells. There is a need for robust, reproducible methods to study single cells with reasonable throughput, as well as to test hypotheses generated by sequencing. Moreover, if these alternative methods were sufficiently accessible, they may be useful in stem cell and developmental biology as a biomarker of ‘stem-ness’, as nuclei tend to compact with differentiation (8–17). However, current methods require fixing cells, stable expression of fluorescent proteins, inaccessible instruments, analysis of subtle changes in heterochromatin morphology, are very low throughput, or are only applicable to certain cell types (see Table 1) (1–8, 15, 18–38), hence current methods have not been widely adopted. In this study, we show that the fluorescence lifetime of SiR-DNA (a far-red nuclear staining dye previously named SiR-Hoechst) (39) can be used as a robust measure of chromatin compaction in live cells, and can be applied to any sample as simply as staining with common live-cell DNA stains.

**Table 1.** A comparison of various methodologies available to assess chromatin compaction. FRET: Förster Resonance Energy Transfer, FRAP: Fluorescence Recovery after Photobleaching, FISH: Fluorescence In Situ Hybridization, ESI: Electron Spectroscopic Imaging, ChIP: Chromatin immunoprecipitation, AFM: Atomic Force Microscopy, Seq: Next Generation Sequencing. *Could provide single gene locus information in combination with dCas9-FP or TALEN-FP, # Throughput is faster with new FLIM systems, $ Single cell Hi-C is possible, but is technically very challenging and expensive. A legend for A,B,C,D,E rankings is included at the bottom with a subjective assessment of instrument availability, throughput, analysis robustness and cell applicability.

| Techniques | References | Gene locus | Single cell | Live cell | Instrument availability | Throughput | Robust analysis | Cell applicability |
| --- | --- | --- | --- | --- | --- | --- | --- | --- |
| SiR-DNA FLIM | This study | No* | Yes | Yes | B | B# | B | A |
| DAPI/Hoechst FLIM | (36, 37) | No* | Yes | No | C | B# | B | A |
| Histone tagged FLIM-FRET sensor | (18–20, 38) | No* | Yes | Yes | B | B# | B | C |
| FRAP | (8, 21) | No* | Yes | Yes | B | C | C | B |
| Histone modification immunofluorescence, FISH | (22–24) | Yes | Yes | No | A | A | E | A |
| Morphology of DAPI or Hoechst staining | (8, 24) | No* | Yes | Yes | A | A | E | E |
| ESI | (25, 26) | No | Yes | No | D | E | A | A |
| ChromEMT | (27) | No | Yes | No | C | E | E | A |
| Chromosome Conformation Capture (3C, 5C, Hi-C) | (1–4) | Yes | No \$ | No | Seq. | Seq. | Seq. | A |
| ChIP, DamID, ATAC-seq, Bisulphite Sequencing | (5–7, 15, 28, 29) | Yes | No | No | Seq. | Seq. | Seq. | A |
| AFM | (30–32) | No | Yes | Yes | B | E | B | D |
| Micropipette squeezing, microfluidic squeezing, substrate stretching | (33–35) | No | Yes | Yes | E | D | D | D |

**Table 1.** A comparison of various methodologies available to assess chromatin compaction.
| Instrument availability | Throughput | Robust analysis | Cell applicability |
| --- | --- | --- | --- |
| A: in most departments, B: in most universities, C: in most countries, D: a few in the world, E: custom-built, Seq.: in most universities | A: 1-10 cells per 10 seconds (confocal z-stack), B: 1-20 cells per 100 seconds (FLIM), C: 1 cell per minute, D: 1 cell per 10 minutes, E: 1 cell per hour, Seq.: One experiment per week | A: direct read-out, B: curve fitting, C: complex curve fitting, D: automated morphology analysis, E: subjective morphology analysis, Seq.: thorough bioinformatics | A: Any cell or tissue, B: cell line transiently expressing transgene, C: clonal cell line stably expressing transgene, D: only isolated nuclei or cells with thin cytosol, E: only cells with a certain nuclear architecture. |

The fluorescence lifetime is the average time a fluorophore spends in the excited state after excitation, and this can be highly sensitive to the fluorophore’s molecular microenvironment (40). Many successful fluorescent biosensors are based on fluorescence lifetime measurements (41–43), as the fluorescence lifetime is generally independent of fluorophore concentration, illumination conditions or microscope setup. The fluorescence lifetime of fluorophores can be mapped spatially using fluorescence lifetime imaging microscopy (FLIM) (44, 45), a technique accessible in many academic microscopy facilities. Throughout this study, the fluorescence lifetime of SiR-DNA was measured using time-correlated single photon counting (TCSPC) FLIM, which provides single-photon sensitivity and the highest signal-to-noise ratio among all the FLIM implementations (46).

We applied FLIM of SiR-DNA dye to easily measure the chromatin compaction in live cells using cell lines treated with artificial compaction or decompaction stimuli, and Embryonic Stem (ES) cells lacking the pluripotency factor NANOG. In addition, we showed that naïve stem cells undergo chromatin decompaction together with Rex1 downregulation as they transition out of the naïve pluripotent stem cell state. Moreover, this technique also works in standard fixed tissue, as we could observe the changes in chromatin compaction in oesophagus epithelium in situ as basal cells differentiate. Therefore, measuring the fluorescence lifetime of SiR-DNA represents an ideally suited method to understand the chromatin compaction state of live or fixed cells with direct relevance to the study of nuclear architecture, stem cells, and cell differentiation.

### Results

To first validate the SiR-DNA FLIM sensor, human fibroblasts and neuroblastoma cells were subjected to treatments that induce chromatin compaction or decompaction (Fig 1). We used ATP starvation (with sodium azide, NaN_3_, and 2-deoxyglucose, 2-DG) to cause chromatin compaction, and histone deacetylase (HDAC) inhibition (with Trichostatin A) to induce chromatin decompaction (18, 19, 36). Cells were subjected to the different treatments and stained with 1 μM SiR-DNA and 10 μM verapamil (a pump inhibitor that improves live cell staining (39)) for one hour, before changing to fresh media lacking SiR-DNA and imaging by FLIM (Fig S1). ATP starvation reduced SiR-DNA fluorescence lifetime and HDAC inhibition increased SiR-DNA fluorescence lifetime significantly, demonstrating that this sensor can measure changes in chromatin compaction status (Fig 1a, b).

**Fig. 1.**
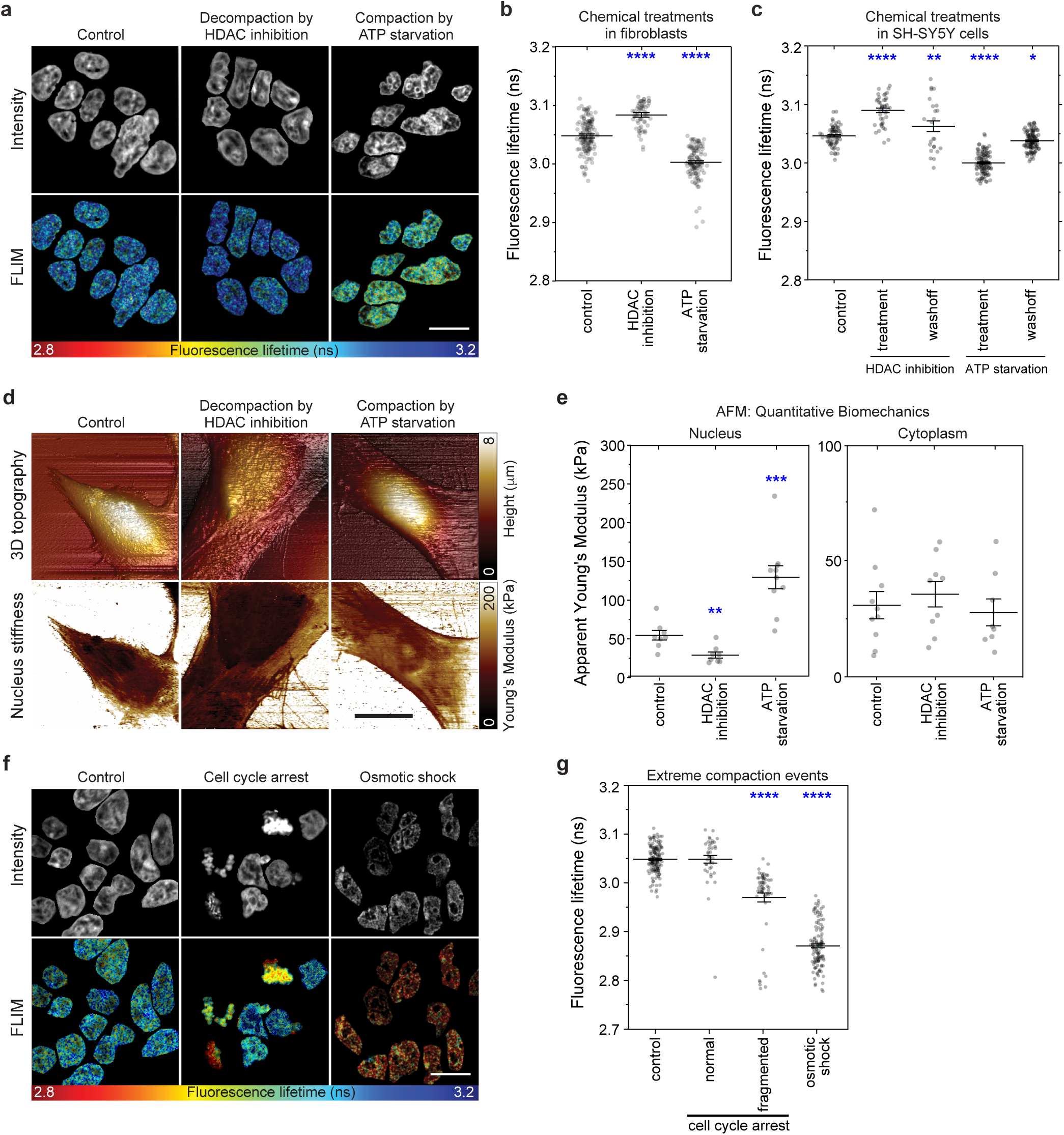
Fluorescence lifetime of SiR-DNA is a novel measure of nuclear compaction in live cells. (a) Representative fluorescence intensity and lifetime images and plot of SiR-DNA fluorescence lifetimes show that treatments causing nuclear compaction such as NaN3+2-DG (ATP starvation) decrease SiR-DNA fluorescence lifetime, whereas treatments causing nuclear decompaction such as Trichostatin A (HDAC inhibition) increase SiR-DNA fluorescence lifetime. Fibroblasts were treated for 1 hour before imaging. 4 independent experiments were performed with >10 images per sample. (c) Similar fluorescence lifetime changes were seen in SH-SY5Y cells undergoing the same treatments. SiR-DNA fluorescence lifetime recovers towards control levels when cells were allowed to recover for 1 hour in untreated media (washoff) before imaging. 3 independent experiments were performed with >10 images per sample. (d) Representative AFM height maps and Young’s Modulus maps and (e) apparent Young’s modulus of nucleus and cytoplasm reveal that compaction or decompaction treatments render nuclei stiffer or softer, respectively, but have no effect on cytoplasm stiffness. Fibroblasts treated as in Fig 1a-b. Each point represents one cell, >3 independent experiments with >3 cells per sample analysed. (f) Representative fluorescence intensity and lifetime images of fibroblasts and (g) plot of SiR-DNA fluorescence lifetimes show that treatments causing extreme nuclear compaction decrease SiR-DNA fluorescence lifetime. Epothilone B (cell cycle arrest) caused some nuclei to adopt a fragmented morphology and only those dying cells had a lower SiR-DNA fluorescence lifetime, and 20% dextrose (osmotic shock) caused a dramatic reduction in SiR-DNA fluorescence lifetime. 4 independent experiments with >10 images per sample analysed. Mean +/- SEM is shown for each plot, where individual points represent one nucleus. Data in b, c, and g were analyzed using unpaired one-way ANOVA, Sidak’s multiple comparisons test where all treatments were compared to control. Data in e were analysed using unpaired, two-tailed t-test with Welch’s correction instead of ANOVA as there are large differences in variances between different treatments. The asterisks in the plots represent significant differences from control: *p<0.05, **p<0.01, ****p<0.0001. All scale bars are 20μm.

SiR-DNA FLIM revealed similar fluorescence lifetime changes in SH-SY5Y neuroblastoma cells subjected to ATP starvation and HDAC inhibition, indicating that these effects are cell type independent (Fig 1c). Moreover, these changes were reversible, as the fluorescence lifetime recovered to untreated cell levels one hour after washing off the various treatments (Fig 1c). The measured compaction (ATP starvation) and decompaction (HDAC inhibition) of chromatin was also confirmed by stiffening and softening of the nucleus, respectively (Fig 1d,e). This was measured using atomic force microscopy (AFM), which has been previously demonstrated to measure nuclear compaction in sufficiently thin cells such as fibroblasts (30–32), but provides very low throughput compared to the SiR-DNA based method. In addition to these relatively mild changes in chromatin compaction, SiR-DNA lifetime showed decreased lifetime in cells dying from cell cycle arrest (epothilone B) and cells undergoing osmotic shock with dextrose, a non-physiological treatment that compacts nuclei by removing water (Fig 1e, f). Notably, the changes in SiR-DNA fluorescence lifetime cannot be attributed to changes in SiR-DNA staining or intensity (Fig S2).

Cutting edge genomics methods such as Hi-C have mapped chromatin interactions during ES cell differentiation in high resolution, however how different ES cell culture conditions affect chromatin compaction and pluripotency remains unclear (reviewed in (47)). Most imaging assays of chromatin compaction have failed to perform well in live cells or required genetic modifications, precluding their adoption in the stem cell field. Stem cell researchers have previously resorted to imaging the chromocentre morphology in fixed cells (8, 24) or less accessible methods such as microfluidic cell squeezing (32) or Electron Spectroscopic Imaging (ESI) (25, 26) (see Table 1). We therefore investigated whether SiR-DNA fluorescence lifetime could provide a quick, affordable, and sensitive method to detect changes in chromatin architecture in live mouse ES cells.

ES cells are characterised by a strikingly open chromatin configuration, including inside constitutive heterochromatin domains (8, 9, 48, 49). The transition from the pluripotent to the committed state features extensive genome reorganisation associated with chromatin compaction and the formation of condensed heterochromatin domains, which is thought to form a repressive environment to facilitate lineage commitment (8–15). Furthermore, differentiating ES cells show a significant increase in cellular (50) and nuclear stiffness (51). Compared to ES cells in serum-LIF medium, ES cells reprogrammed to a naïve state using 2i-LIF medium (serum-free N2B27 medium containing inhibitors of the MEK and GSK3*β* pathways) (52, 53) are characterised by higher cellular homogeneity and low levels of DNA methylation, which has been strongly associated with a more permissive chromatin state (29, 47, 54–56). Surprisingly, despite the lower DNA methylation, the SiR-DNA fluorescence lifetime was reduced in 2i-LIF cultured ES cells, indicating that the chromatin in these naïve cells was significantly more compact compared to ES cells maintained in serum-containing medium (Fig 2a, b). To verify our observation that the chromatin was compacted in these ES cells using an established assay, we also examined the chromocenter morphology (8, 24). Naïve pluripotent ES cells in 2i-LIF medium showed an increase in chromocenter numbers (Fig S3), confirming a general compaction of the genome during reversion to the naïve pluripotent state. Our data support that naïve ES cells transition through a less-compact ‘formative period’ (57) state before differentiating and compacting. Indeed, we found that in a time-course experiment, as naïve Rex1-GFP-expressing cells left the naïve state and downregulated Rex1-GFP (58), they had an increased SiR-DNA fluorescence lifetime indicating chromatin decompaction (Fig 2c, S4).

**Fig. 2.**
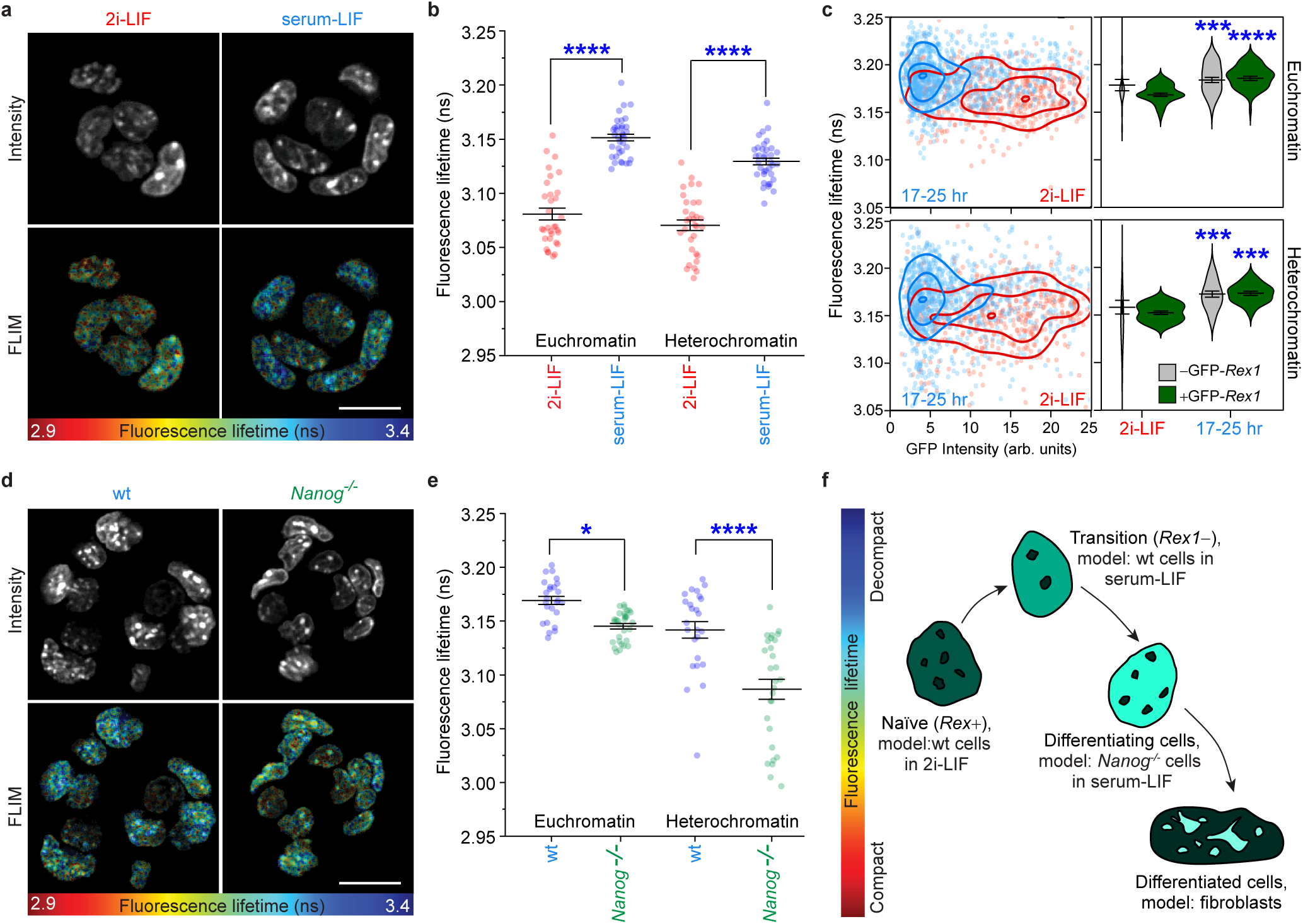
SiR-DNA fluorescence lifetime analysis reveals that murine naïve embryonic stem cells transit through a genome decompaction stage before committing to differentiation. (a) Fluorescence intensity and lifetime images and (b) plot of SiR-DNA fluorescence lifetimes show that naïve ES cell nuclei (2i-LIF medium) display a lower fluorescence lifetime (more compact) than transition ES cell nuclei (serum-LIF medium), in both euchromatin and heterochromatin regions. Each point represents one image, 3 independent experiments with >8 images per sample. Unpaired one-way ANOVA with Sidak’s multiple comparisons test. (c) Rex1-GFP ES cell time course after 2i withdrawal shows that nuclei increase their SiR-DNA fluorescence lifetime (become less compact) as they lose Rex1-GFP expression (exit naïve state). Each point represents one cell, 3 independent experiments with >9 images per sample. Threshold for ‘GFP-positive’ is 5 intensity units. All timepoints are compared against the GFP positive 2i-LIF condition using unpaired one-way ANOVA with Dunnett’s multiple comparisons test. (d) Fluorescence intensity and lifetime images and (e) plot of SiR-DNA fluorescence lifetimes show that *Nanog-/-* cell nuclei display a lower fluorescence lifetime (more compact) than wildtype ES cell nuclei (both in serum-LIF), in both euchromatin and heterochromatin regions. Each point represents one image, 3 independent experiments with >8 images per sample analysed. Unpaired one-way ANOVA with Sidak’s multiple comparisons test. (f) Model of ES cell differentiation, showing that naïve ES cells (Rex1 positive cells in 2i-LIF) first go through a chromatin decompaction stage (Rex1 negative cells in serum-LIF), before further differentiation and chromatin compaction (*Nanog-/-* cells, fibroblasts, and SH-SY5Y cells). Mean +/- SEM is shown for each plot. * p<0.05, ** p<0.01, *** p<0.001, **** p<0.0001. Scale bars are 30μm.

Once through the formative transition state, ES cells start to differentiate and the chromatin starts to compact (8–15). The pluripotent transcription factor NANOG is required to maintain a globally open chromatin in ES cells as *Nanog-/-* nuclei were shown to be more compact using ESI and scoring changes to chromocenter morphology (8, 24). In this study, we have confirmed this finding as *Nanog-/-* ES cell nuclei displayed a shorter SiR-DNA fluorescence lifetime compared to wild-type cells (Fig 2d,e). With these data we can draw a simplistic model encompassing the spectrum of chromatin compaction and measurable by FLIM of SiR-DNA (Fig 2f), where naïve ES cells (represented in this study by ES cells in 2i-LIF) first decompact (represented by ES cells in serum-LIF or 17-25 hours after 2i withdrawal), and then start to compact (represented by *Nanog-/-* ES cells), and subsequently differentiate into the various mature cell types with very compact chromatin (represented by fibroblasts and SH-SY5Y cells).

While the ability to measure chromatin compaction in live cells may be optimal for cells in culture, we also found that SiR-DNA fluorescence lifetime can be applied to fixed tissues allowing the study of chromatin architecture *in situ* with no genetic modification required. Mouse oesophagus cryosections were stained with SiR-DNA as well as wheat-germ agglutinin (WGA) to label epithelial cell membranes and imaged with FLIM (Fig 3a,b). The basal cells (stem cells) had a higher fluorescence lifetime than the suprabasal (differentiating) cells, which is expected since extensive genome reorganisation associated with chromatin compaction occurs upon differentiation (59). Interestingly, there was a large variation in the fluorescence lifetimes of the suprabasal cells, including many nuclei containing a high-lifetime spot. This suggests that there may be a general compaction of the chromatin during differentiation, coupled with a partial decompaction of some regions of the nucleus involved in the transcription of keratin and other keratinocyte genes (59). These data demonstrate that measuring SiR-DNA fluorescence lifetime is an effective new tool that can contribute to studies of nuclear architecture and stem cell differentiation *in situ*, in ex-vivo tissue sections.

**Fig. 3.**
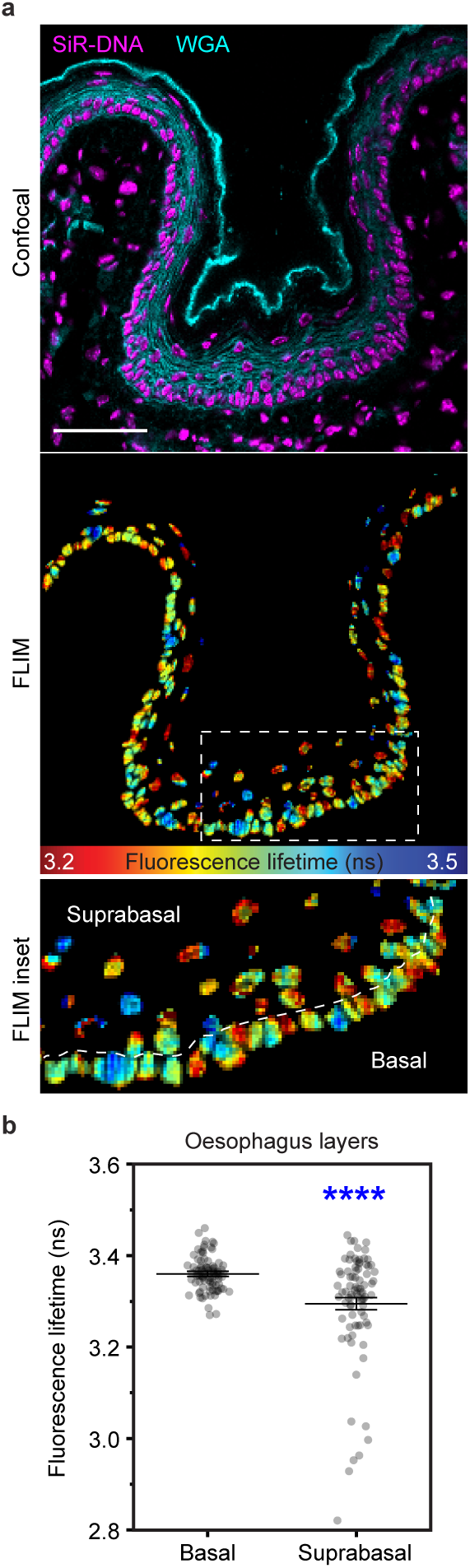
Chromatin compaction in fixed tissues. (a) Fluorescence intensity and lifetime image and (b) plot of SiR-DNA fluorescence lifetimes in fixed mouse oesophagus tissue show a longer lifetime (less compact) in basal stem cells than differentiating suprabasal cells. Wheat Germ Agglutinin (WGA) labels epithelial cell membranes. Each point represents one nucleus, data from one representative image shown, out of 6 mice analysed. Unpaired t-test with Welch’s correction. Mean +/- SEM is shown, **** p<0.0001. Scale bar: 50μm.

## Discussion

In this study, we have shown that SiR-DNA FLIM is an accurate and quantitative measure of chromatin compaction. FLIM of SiR-DNA provides excellent sensitivity and allows for imaging any cell in their native state without requiring fluorescent protein expression. While FLIM is not a very common microscopy technique, there are FLIM systems available in the microscopy suites of many academic institutes, and recent developments in FLIM technology are making FLIM easier, faster and more accessible than ever.

First, we validated the assay in fibroblasts and SH-SY5Y cells using standard chemical treatments to compact and decompact chromatin. We then illuminated new details of chromatin decompaction during naïve ES cell differentiation, as well as confirming recently published results on the nuclear architecture of ES cell knockouts of Nanog. Finally, we showed that this method is amenable to studies of chromatin compaction in fixed tissue slices, demonstrating that this method really is applicable across biological contexts with no genetic modification required. We have shown that the measurement of SiR-DNA fluorescence lifetime is compatible with expression of other fluorescent markers, such as Rex1-GFP to mark naïve ES cells or WGA staining to mark epithelial cells. We expect that this method could also be used in combination with fluorescent dCas9 or fluorescence *in situ* hybridisation (FISH) to examine the local chromatin state of particular loci.

Our data highlight the need for independent methods to measure chromatin compaction, as our observations challenge previous assumptions that lower levels of DNA methylation in 2i-treated ES cells (29) also results in less compaction. There have been some reports suggesting that the epigenetic marker H3K27me3 may be able to repress parts of the genome and counterbalance lower levels of DNA methylation, at least within some loci of 2i-treated cells (60, 61).

Our results showing that naïve ES cells decompact upon 2i withdrawal support previous data that Rex1 downregulation occurs at the same time as a transition to an auxetic nucleus phenotype (32). Therefore, this decompacted chromatin state may be part of the same formative period (57) whereby naïve ES cells must first decompact to reach a transition state where chromatin is accessible to a gene regulatory switch to gain competence for lineage commitment (14, 21, 32, 62).

Chromatin compaction has been shown to correlate with differentiation state, as differentiated cells generally compact the regions of their genome not required for their mature function to maintain their identity (8–17). In this study we have shown some notable exceptions to this rule, as naïve ES cells initially decompacted upon leaving the naïve state, and differentiating keratinocytes appear to have foci of decompaction, but as long as these types of exceptions are kept in mind, this method may be viable as a general ‘stem-ness’ assay. Moreover, we foresee that this method could be amenable to a plate-reader format for high content screens of stem cell phenotypes.

## ACKNOWLEDGEMENTS

This project has received funding from the European Union’s Horizon 2020 research and innovation programme under Grant Agreement No. 722380 (SUPUVIR). G.S.K.S. and C.F.K. acknowledge funding from the Wellcome Trust, the UK Medical Research Council (MRC), Alzheimer Research UK (ARUK), and Infinitus China Ltd. C.F.K. acknowledges funding from the UK Engineering and Physical Sciences Research Council (EPSRC). P.J.R.-G. acknowledges funding from the UK Biotechnology and Biological Sciences Research Council (BBSRC) (BB/M022285/1 and BBS/E/B/000C0422) and the European Commission Network of Excellence EpiGeneSys (HEALTH-F4-2010-257082). D.F-A. was supported by a core grant from the Wellcome Trust to the Wellcome Sanger Institute. We thank Ricardo Henriques lab for this LaTeX template.

## AUTHOR CONTRIBUTIONS

C.H. and C.P. performed the FLIM imaging and analysis. C.H. and K.A.F. prepared fibroblast and SH-SY5Y samples. I.M. performed the AFM studies and analysis. D.F-A. prepared oesophagus tissue samples. C.L.N. and P.J.R-G. prepared ES cell samples and performed chromocenter analysis. M.S.H and K.C. prepared Rex1-GFP ES cell samples. P.P.V wrote scripts for image analysis. C.H., C.P., K.A.F, C.F.K., and G.S.K.S. conceived and designed the experiments. C.H. and C.P. wrote the manuscript with help from C.N. and G.S.K.S.

## COMPETING FINANCIAL INTERESTS

The authors declare no competing financial interests.

## Supplementary Note 1: Materials and Methods

### Cell culture

Human HFF-1 fibroblasts from the American Type Culture Collection were cultured in Dulbecco’s Modified Eagle’s Medium (Gibco, Thermo Fisher Scientific, Waltham, MA, USA) with 10% fetal bovine serum (FBS, Gibco), 1% GlutaMAX (Gibco), and 1% antibiotic-antimycotic (Gibco). Human SH-SY5Y neuroblastoma cells from the European Collection of Cell Cultures were cultured in 41% Minimum Essential Medium (Gibco), 41% Ham’s F-12 Nutrient Mixture (Gibco), 15% FBS, 1% non-essential medium (Gibco), 1% GlutaMAX (Gibco), 1% antibiotic-antimycotic (Gibco).

Mouse E14Tg2a (also referred to as E14) is a male mESC line of 129/Ola background (1). E14Tg2a, E14Tg2a-derived *Nanog*^*-/-*^ (RCNβH-B(t)) (2), were cultured on gelatine-coated plastic in standard ESC medium (DMEM with 15% FBS, 1 mM sodium pyruvate, 0.1 mM 2-mercaptoethanol, 0.1 mM nonessential amino acids, 1% GlutaMAX, 1000 U/mL LIF). For naïve reprogramming, E14 cells and Rex1GFPd2 reporter mESCs were cultured on gelatine-coated plastic (coated with 0.1% gelatine in PBS, RT for 30 minutes) in serum-free N2B27 supplemented with 2i inhibitors, 1uM PD0325901 (MEK inhibitor) + 3 μM Chiron (CHIR99021, GSK3 inhibitor) + 100U/ml LIF. N2B27 was made with equal parts of Neurobasal media (Gibco) and Invitrogen DMEM F-12 (Gibco), supplemented with 0.1 mM 2-mercaptoethanol (ThermoFisher), 0.5% N2 (made in-house (3)), 1% B27 (ThermoFisher), 2mM L-glutamine (ThermoFisher), 0.1% BSA (ThermoFisher) and 12.5 μg/ml human recombinant insulin zinc (ThermoFisher). mESCs cultured in 2i were split with Accutase (Millipore) every 2-3 days and seeded at 1.5×10^4^ cells/cm^2^. Exit from naïve pluripotency was initiated by re-plating cells in N2B27 without 2i inhibitors at a density of 1×10^4^ cells/cm^2^ on laminin coated dishes (coated overnight incubation at 10 μg/ml at 4°C) at 3, 9, 17, 25, 34, and 48 hours before imaging.

### SiR-DNA staining

For SiR-DNA-based analysis of chromatin compaction by FLIM, cells were plated in 35 mm glass bottom dishes (P35-1.5-14-C, Mattek Corporation, MA, USA) the day before the start of the experiment. Cells were stained with 1 μM SiR-DNA (Spirochrome Ltd., Stein am Rhein, Switzerland) with 10 μM verapamil (Spirochrome Ltd.) in cell culture medium for 1 hour, before changing to cell culture medium containing 10 μM verapamil and 20 mM HEPES pH 7.4 for imaging. Nuclear compaction or decompaction treatments (10 mM sodium azide and 50 mM 2-deoxyglucose; 200 ng/ml Trichostatin A; or 20% dextrose) were included in the staining and imaging medium.

### Mouse Oesophagus

Optimal cutting temperature compound (OCT) embedded C57BL/6J oesophageal cryosections of 10μm thickness were fixed with 2% paraformaldehyde for 5 min. After two washes in PBS, slides were incubated in blocking buffer (0.5% bovine serum albumin, 0.25% fish skin gelatine, 0.5% Triton X-100 and 10% donkey serum, in PBS) and stained in the same buffer with SiR-DNA and Alexa Fluor 488 Wheat Germ Agglutinin for 5 minutes at room temperature. Samples were washed twice in PBS and mounted using 50/50 PBS glycerol.

### Fluorescence lifetime imaging (FLIM)

All samples were assayed on a home-built confocal platform (Olympus Fluoview FV300) integrated with time-correlated single photon counting (TCSPC) to measure fluorescence lifetime in every image pixel. To excite SiR-DNA, output from a pulsed supercontinuum source (WL-SC-400-15, Fianium Ltd., UK, repetition rate 40MHz) was filtered using an acousto-optic tunable filter (AA Optoelectronic AOTFnC-VIS) to obtain a narrow wavelength band centered at 640nm and an additional bandpass filter FF01-635/18 was used to cleanup excitation light. Fluorescence emission from the sample was filtered using 700/70nm (Comar Optics, U.K.) before passing onto a photomultiplier tube (PMC-100, Becker&Hickl GmbH, Berlin, Germany). Photons were recorded in time-tagged, time-resolved mode that permits sorting photons from each pixel into a histogram according to their arrival times. The data was recorded by a TCSPC module (SPC-830, Becker&Hickl GmBH). Photons were acquired for 200 seconds to make a single 256 × 256 FLIM image. Photon count rates were always kept below 1% of the laser repetition rate to prevent pulse pile-up. Photobleaching was verified to be negligible during acquisition. Approximately 10 representative images with several nuclei per field of view were acquired for each condition.

### FLIM Analysis

All FLIM images were analysed using FLIMfit v4.12.1(4) and fitted with a monoexponential decay function (S1). For the analysis of fibroblasts and SH-SY5Y cells, an average fluorescence lifetime per nucleus was obtained from a large field of view using intensity-based segmentation. For images displayed in figures, lifetimes were fitted per pixel. For the analysis of stem cell nuclei images in 2, whole nuclei were segmented based on intensity (debris and mitotic nuclei were manually removed). A two-level mask separating heterochromatin spots and euchromatin (S5) was created with Icy software spot detection tool (5, 6). Pixels in these two chromatin regions were binned and fitted separately to obtain two lifetime values for each image (demonstrated in Fig 2). For the analysis of Rex1-GFP stem cell nuclei, the masks were separated into single cells using a custom Matlab (MathWorks, MA, USA) script, to obtain the lifetime of the euchromatin and heterochromatin in each cell.

These masks were also used to measure the corresponding GFP intensity of each cell, and the lifetime and GFP intensity data were combined (Fig S4) using R (7, 8). Data were plotted with Origin 2018b (OriginLab, Northampton, MA) and statistical analysis was carried out using Graphpad Prism 7 (La Jolla, California, USA).

### Atomic Force Microscopy

For AFM nuclear stiffness measurements, cells were plated at 2×10^5^ cells per 50 mm glass bottom dish (GWST-5040, Willco Wells BV, Amsterdam, The Netherlands) the day before imaging. Cells were treated with nuclear compaction or decompaction treatments (see above) in cell culture medium for 1 hour, before changing to Dynamic Imaging Medium (150 mM NaCl, 5 mM KCl, 1 mM MgCl_2_, 1 mM CaCl_2_, 5 mM glucose, 10 mM HEPES pH 7.4) containing the chemical treatments during AFM imaging.

Atomic force microscopy measurements were performed on a Bioscope Resolve bioAFM (Bruker, Billerica, MA, USA), operated in PeakForce QNM mode. Live Cell probes (PFQNM-LC, Bruker AFM probes) were used for all experiments. The probes were pre-calibrated for spring constant (nominal 0.08 N/m) and its deflection sensitivity was measured at the start of the experiment, using a no-touch calibration. The sample stage was heated and maintained at 37°C. The force applied to the cells was kept constant throughout the experiments and was less than 1 nN, with typical values being 400-600 pN.

### Analysis of Chromocenter Morphology

DAPI linescan analyses were performed using ImageJ on optical sections where DAPI foci were at optimal focal planes. Fluorescence intensity histograms were generated through the nucleus and the background (outside the nucleus) was subtracted from the nucleoplasmic (baseline fluorescence within nucleus) and chromocentre signals. Variations in these data were calculated as a ratio of chromocentre peak height to nucleoplasmic signal.

---

**Fig. S1.**
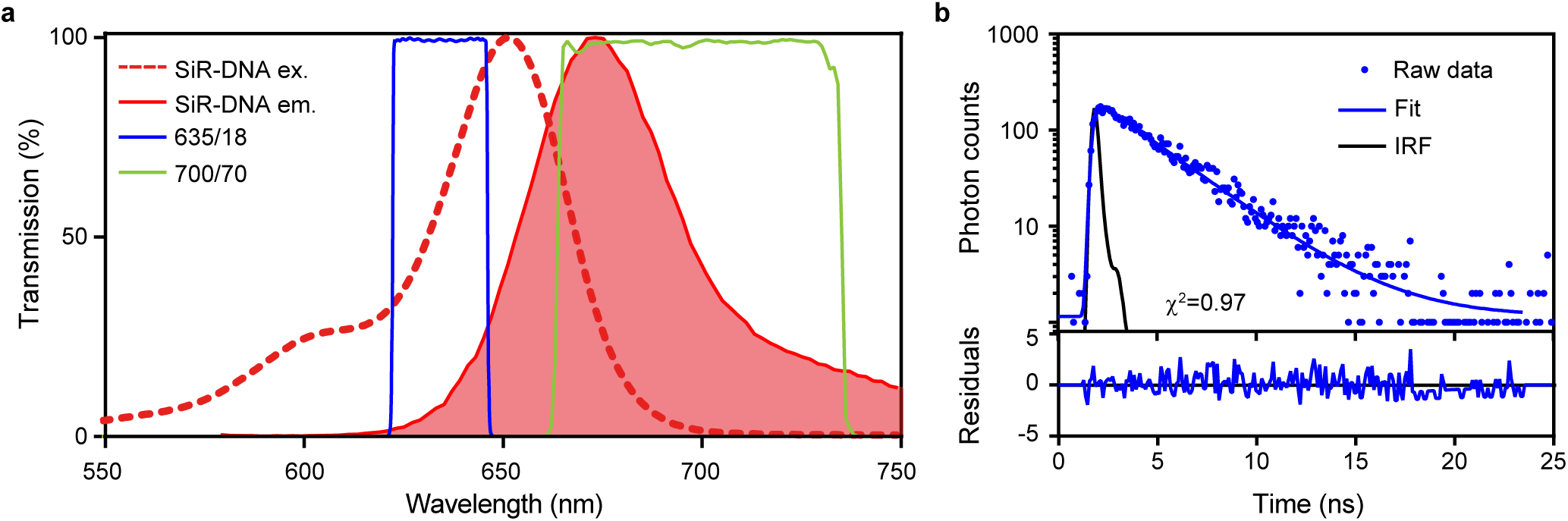
Fluorescence properties of SiR-DNA. (A) Excitation and emission spectra from the Spirochrome Ltd., with excitation and emission filters used in this study. (B) Typical fluorescence decay curve of SiR-DNA from a 3*×*3 pixel bin, with a monoexponential decay fitted using FLIMfit and fit residuals (error) plotted. The fluorescence decay curve is from a representative TCSPC-FLIM image of fibroblasts stained with SiR-DNA. IRF: Instrument Response Function

**Fig. S2.**
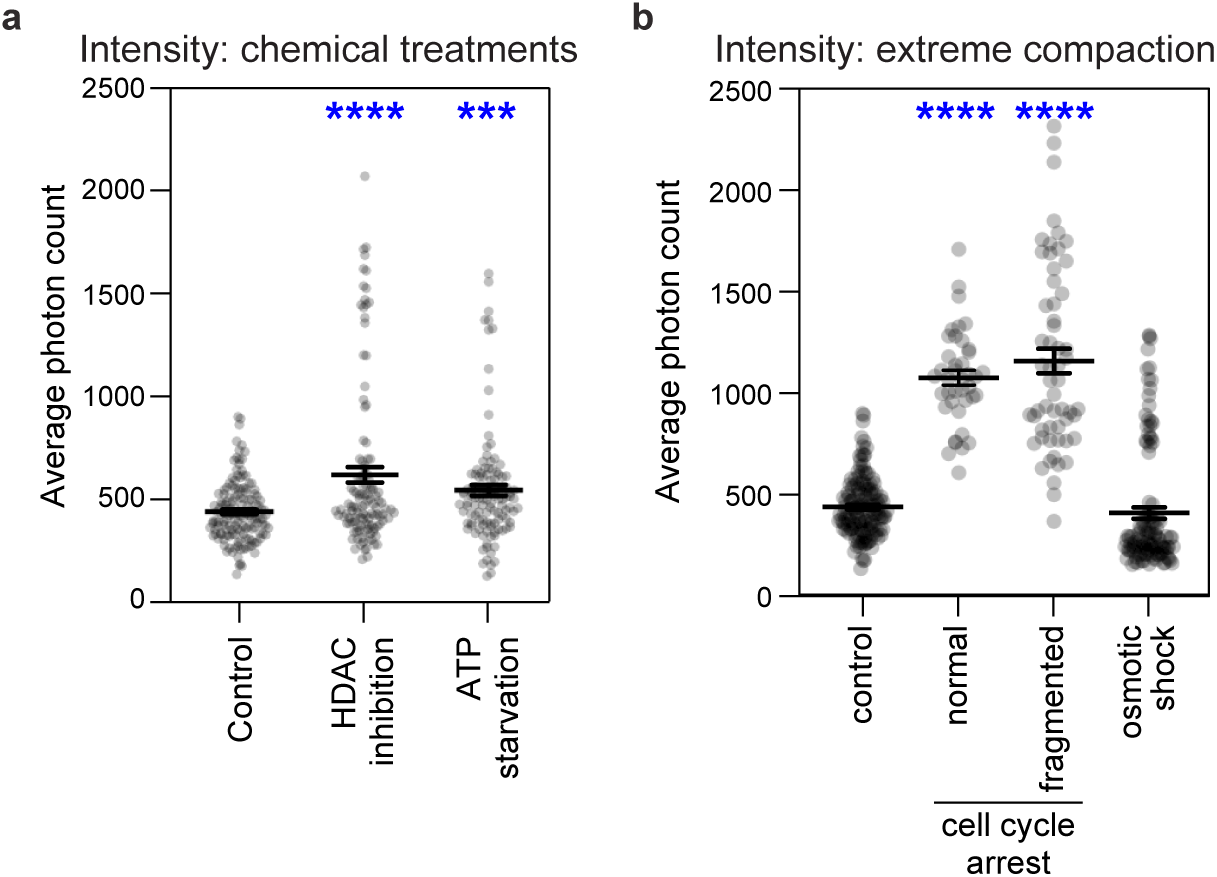
SiR-DNA fluorescence intensity does not correlate with chromatin compaction. (a) Fluorescence intensity from fibroblast images used in Fig 1a and (b) Fluorescence intensity from fibroblast images used in Fig 1e show that different samples have different fluorescence intensities, but these do not correlate with chromatin compaction or decompaction. All treatments compared to control, unpaired one-way ANOVA with Sidak’s multiple comparisons test. Mean +/- SEM is shown for each plot. ***p<0.001, ****p<0.0001.

**Fig. S3.**
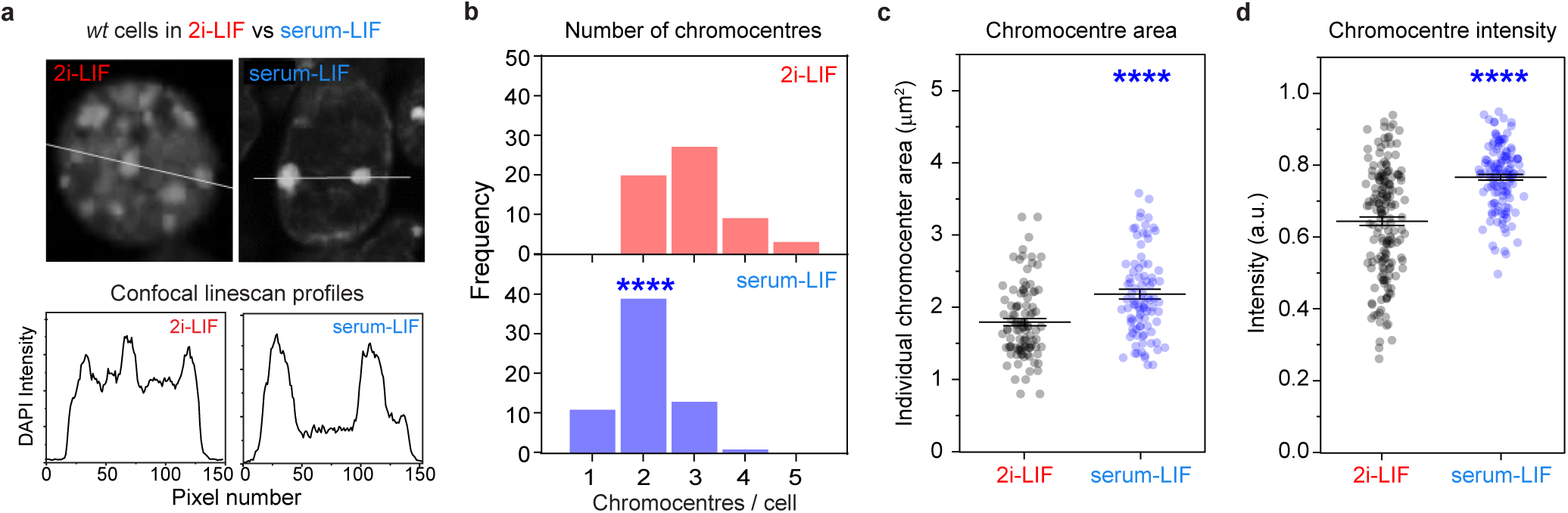
Chromocentre morphology analysis of DAPI-stained ES cells shows that naïve cells (2i-LIF medium) had decreased chromocentre intensity and total chromocentre area compared to transition cells (serum-LIF medium). (a) Linescan profile and (b) chromocenter counting of DAPI stained ES cells show that naïve ES cell nuclei (2i-LIF medium) had increased numbers of chromocenters but (c) reduced area of individual chromocentres. (d) Chromocentre intensity analysis shows that naïve ES cells (2i-LIF medium) also had reduced chromocentre intensity. 3 independent experiments with >40 cells were analysed per sample. Mean +/- SEM is shown for each plot. Data in b, c and d were analysed using unpaired two-tailed t-test with Welch’s correction. ****p<0.0001.

**Fig. S4.**
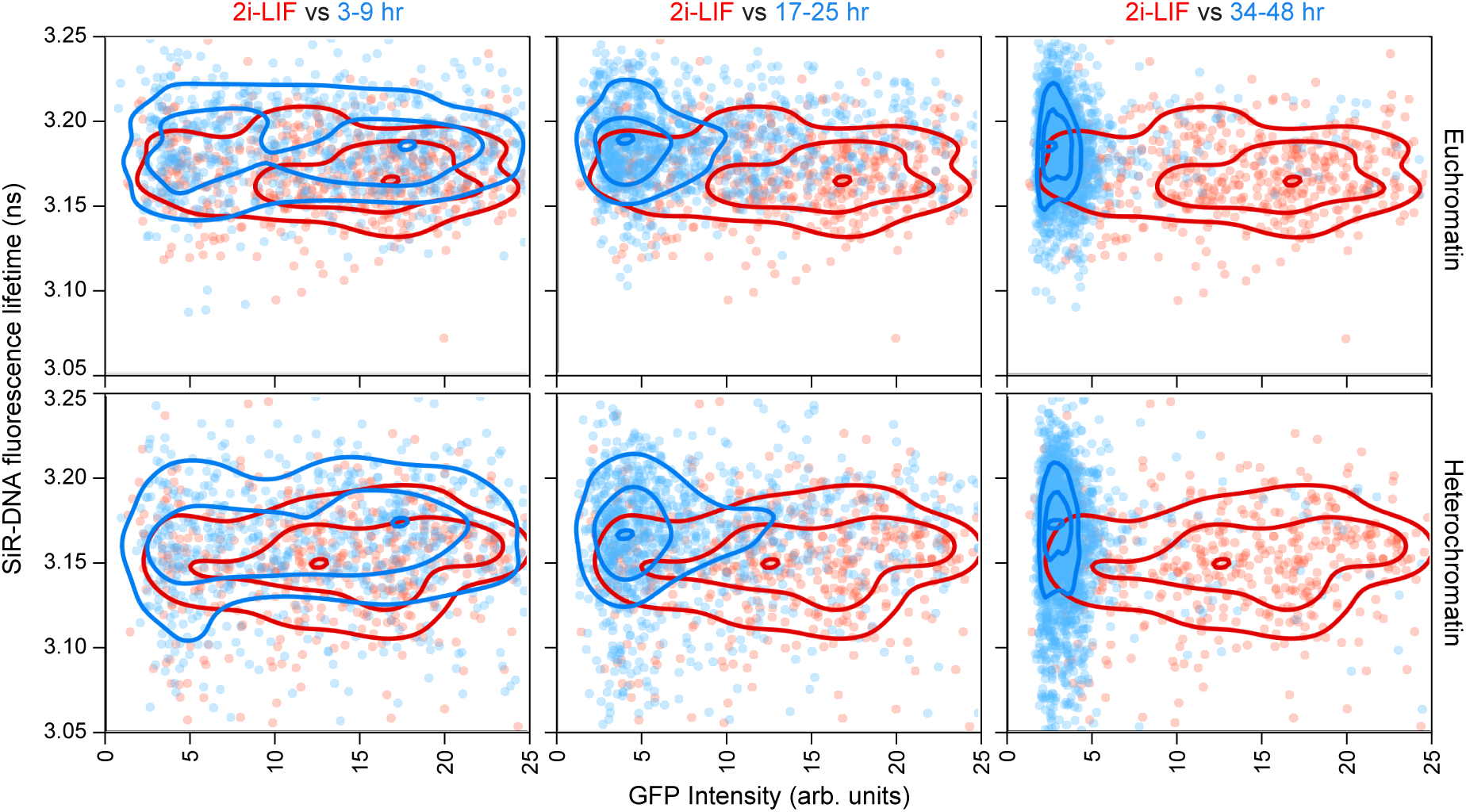
Timecourse of 2i withdrawal in Rex1-GFP ES cells shows that naïve embryonic stem cells have undergone genome decompaction in both euchromatin and heterochromatin. Single-cell analysis of GFP intensity (Rex1-GFP level) and SiR-DNA fluorescence lifetime (chromatin compaction) shows that cells lost Rex1 expression and increased SiR-DNA fluorescence lifetime (decompacted), as they transitioned out of the naïve cell state. Each point represents one cell, 3 independent experiments with >9 images per sample.

**Fig. S5.**
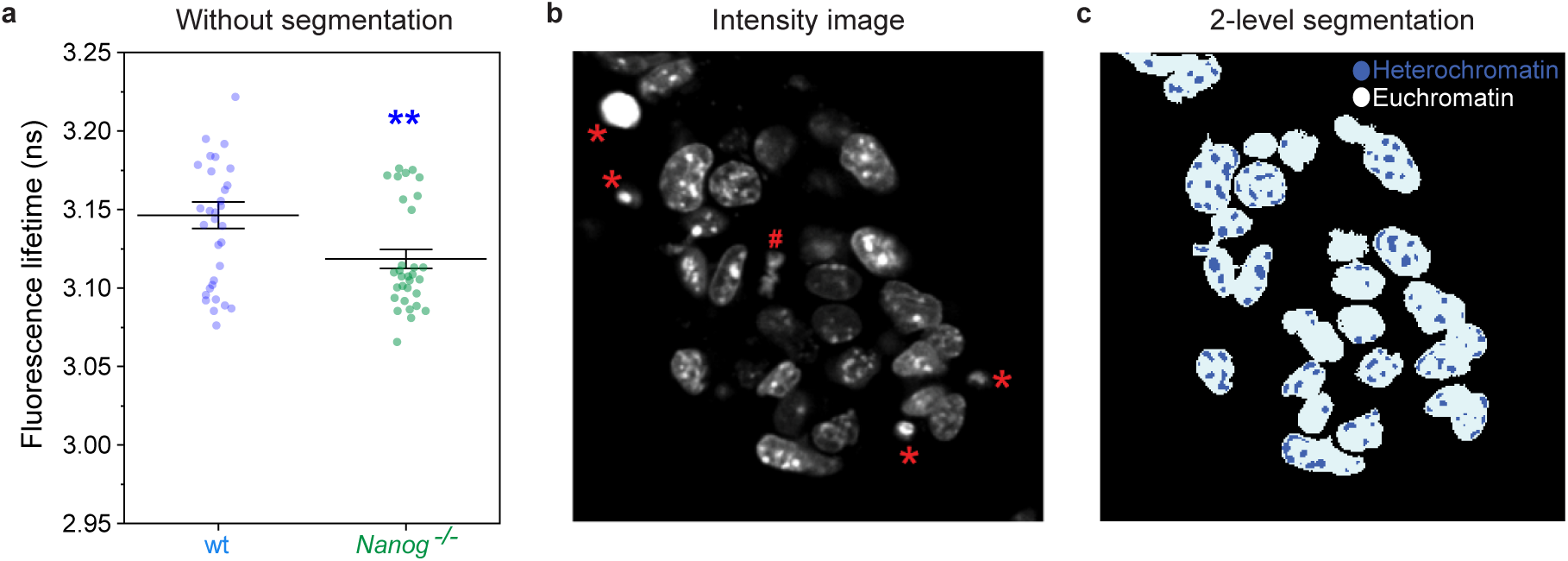
Spot detection strategy to segment euchromatin from heterochromatin. (a) Plot of SiR-DNA fluorescence lifetimes show that Nanog-/- ES cell nuclei have a lower fluorescence lifetime (are more compact) than wildtype ES cell nuclei, even without segmentation of nuclei into heterochromatin and euchromatin. These plots are based on the same data as in Figure 2d and e. Mean +/- SEM is shown in the plot. Unpaired t-test with Welch’s correction, ** p<0.01. (b) Representative intensity image. A threshold was used to segment the nuclei, and then debris (*) and mitotic nuclei (#) were manually removed from the mask. The Spot Detector algorithm using the image analysis software Icy was used to segment the heterochromatin spots. The nuclei mask and the heterochromatin mask were multiplied together to remove spots detected outside the curated nuclei, and the resulting curated heterochromatin mask was added to the nuclei mask to make a two-level mask for import into the FLIMfit Segmentation Manager. (c) Image of the resulting two-level mask, with euchromatin in white and heterochromatin in blue.

## Notes

### Competing Interest Statement

The authors have declared no competing interest.

